# Acute partner loss enhances social motivation and nucleus accumbens dopamine release in prairie voles

**DOI:** 10.64898/2026.09.22.753547

**Authors:** Natsumi Komatsu, Alexis M. Black, Stephen E. Song, Markita P. Landry, Annaliese K. Beery

## Abstract

The loss of close social relationships triggers profound mental and physical health consequences across social species. We utilized the socially selective prairie vole (Microtus ochrogaster) to investigate how 5-day separation from a bonded mate or peer partner shapes social motivation and dopamine release kinetics in males and females. Prior to partner loss, baseline social motivation to access a partner or a novel object differed by sex and relationship type (mate or peer partnership). Across all groups, however, separation induced heightened partner-seeking. Partner loss enhanced dopamine signaling in the nucleus accumbens, increasing evoked dopamine release, the number of putative dopamine release sites, and tyrosine hydroxylase abundance, regardless of sex or relationship type. These findings suggest that mesolimbic reward circuitry rapidly adapts to the loss of a bonded partner, signaling a critical social deficit across relationship frameworks. Furthermore, by leveraging synthetic sensors, this study provides novel subcellular insights into presynaptic dopaminergic plasticity following 5-day relationship disruption, directly linking relationship loss to heightened social motivation and shifts in reward signaling.

**Significance Statement:** From friendships to mate partnerships, reciprocal social connections are vital for humans and other species that form social bonds. This study explores the behavioral and neural consequences of disrupting these relationships using the prairie vole, a species capable of forming selective bonds with both peers and mates. We demonstrate that, despite baseline sex and relationship-type differences in social reward, separation from a bonded partner triggers a uniform enhancement in social drive accompanied by elevated accumbal dopamine release and synthesis. This reveals a conserved, homeostatic reward mechanism that treats the loss of relationships as a critical deficit, advancing our understanding of the neurobiology of social bonding as well as grief and loneliness.

## Introduction

Social relationships are fundamental for mammalian survival, well-being, and longevity. In humans, social connectedness is among the most powerful determinants of health across the lifespan, predicting immune function, stress resilience, and cardiovascular health (1–4). Conversely, loneliness and social isolation have profound detrimental effects on both physical and mental health (5, 6), conferring a risk of mortality comparable to that associated with smoking and obesity (7, 8). The loss of a loved one represents a particularly profound form of social disconnection that can be especially impactful, escalating risks of cardiovascular disease (9, 10), suicide and mortality (11, 12), and psychiatric conditions such as prolonged grief disorder (13, 14).

Prairie voles (*Microtus ochrogaster*) provide a powerful opportunity to investigate the impacts of relationship loss on health and behavior. Unlike traditional laboratory rodents, prairie voles form selective and enduring relationships, including socially monogamous pair bonds with their mate (15–17), as well as friendship-like peer relationships with non-reproductive companions (18–20). These social bonds enable the study of specific partner loss and allow us to evaluate whether the disruption of different relationship types triggers common neural and behavioral responses. Prior work in prairie voles has shown that chronic isolation increases anxiety- and depressive-like behaviors (21, 22). These behavioral changes are accompanied by significant neuroendocrine shifts, including increased circulating oxytocin and corticosterone (23–25), alongside heightened oxytocin, vasopressin, and corticotropin-releasing factor activity in the hypothalamus, evidenced by elevated mRNA expression and immunoreactivity (25, 26). Partner loss in voles also disrupts neuroendocrine and cardiac function, leading to endothelial impairment (27, 28), and triggering increased heart rate, dysregulated cardiac rhythm, and heightened corticosterone levels after only 5-day separation(29). In male voles, 4-day separation from a female mate partner produces more severe anxiety- and depressive-like behaviors than separation from a male peer (30). Peer and mate loss also have different effects on odor preferences in females (31). Thus, relationship type may have effects on social effort and/or neurochemical changes following loss.

The deleterious effects of isolation have been documented across a wide range of social species, from bumblebees to elephants (32, 33), highlighting the widespread importance of social connectivity. This phenomenon has been particularly well-studied in laboratory rodents, which exhibit a well-documented “isolation syndrome” following prolonged separation (2–13 weeks). This condition is characterized by both physiological and behavioral dysfunction, including endocrine changes, weight loss, altered hematology, heightened pain sensitivity, cognitive and reproductive deficits, and increased aggression (34, 35). Isolation is further associated with suppressed neurogenesis (22, 36) and compromised cardiac function (37, 38). Evidence suggests that females are particularly vulnerable to the impacts of isolation compared to males (35, 39), and the severity of these effects varies by the developmental stage at the time of exposure (40). General social deprivation similarly heightens social motivation, social investigation, and affiliative behavior in rats and mice following both chronic (>1 week) and acute (<1 week) social deprivation (41–48). Such increases in social interest have been recently understood in the context of disrupted social homeostasis, where deficits in social interaction lead to social “craving” or “hunger,” driving the increased social seeking that follows social isolation (49). In prairie voles, chronic social isolation increased sociability toward an unfamiliar vole (22), and one week of separation from a bonded mate increased investigation of bedding from their former shared cage (31, 50). In contrast, brief 3-day isolation did not affect affiliative contact during 30 minutes of direct social interaction with a novel same-sex or opposite-sex conspecific, but in some cases increased male aggression (51).

Elevated social seeking after isolation is thought to reflect heightened mesolimbic dopamine signaling, yet the presynaptic adaptations that drive this shift remain largely unresolved. The nucleus accumbens (NAc) receives dense dopaminergic innervation and serves as a critical hub for social reward (52), making it a prime candidate region for linking isolation-induced changes in dopamine dynamics to social motivation. Using fast-scan cyclic voltammetry and microdialysis, prior work has shown that six weeks of chronic, early-life social isolation increases dopamine release in rats (53) and that daily parental separation on postnatal days 1-13 produces a similar increase in mandarin voles (54). Separately, 24-hour isolation potentiates dopamine synapses in the dorsal raphe and elevates their activity during social reunion in mice (55), indicating that even brief separation can rapidly remodel dopaminergic regions that supply the NAc. Yet how acute, adult separation affects dopamine release within the NAc itself has not been examined in any rodent, and no prior study has resolved these dynamics at the level of individual release sites — existing work relies on population-level measurements that cannot capture subcellular changes in release probability or site number. Isolation- and loss-induced dopamine release has also never been measured directly in prairie voles, despite evidence that, in mate-paired voles, seeking and approaching a bonded partner evoke greater accumbal dopamine release than the same behaviors directed at an unfamiliar vole (56). Together, these findings suggest that partner separation could alter dopamine signaling through two non-exclusive routes — the general consequences of social isolation, and the specific loss of a preferred partner.

Thus, the consequences of acute partner loss in adulthood remain poorly understood, particularly across relationship types and in females. Moreover, little is known about how partner separation alters the function of dopaminergic neurons at the subcellular level. The present study asked how 5-day partner separation impacts subsequent social motivation for partner access and dopamine release dynamics in adult prairie voles. Specifically, we evaluated these effects across two axes: relationship type (peer relationships vs. mate relationships) in females, and biological sex (female vs. male) in mate relationships. We show that partner separation increases social motivation and accumbal dopamine release across relationship types and sexes, revealing subcellular dopaminergic adaptations that accompany relationship loss.

## Results

### Partner loss increased partner seeking

All groups worked significantly harder to access their partner versus a novel object following social loss, as shown by a significant post-separation increase in social reward preference score within female peers (*Figure 1A*; t(6)=5.061, *p*=0.0023), female mates (*Figure 1B*; t(8)=3.093, *p*=0.0148) and male mates (*Figure 1C;* t(5)=2.906, *p*=0.0336). The social reward preference score was calculated as the number of partner rewards/(partner+novel object rewards).

**Figure 1.**
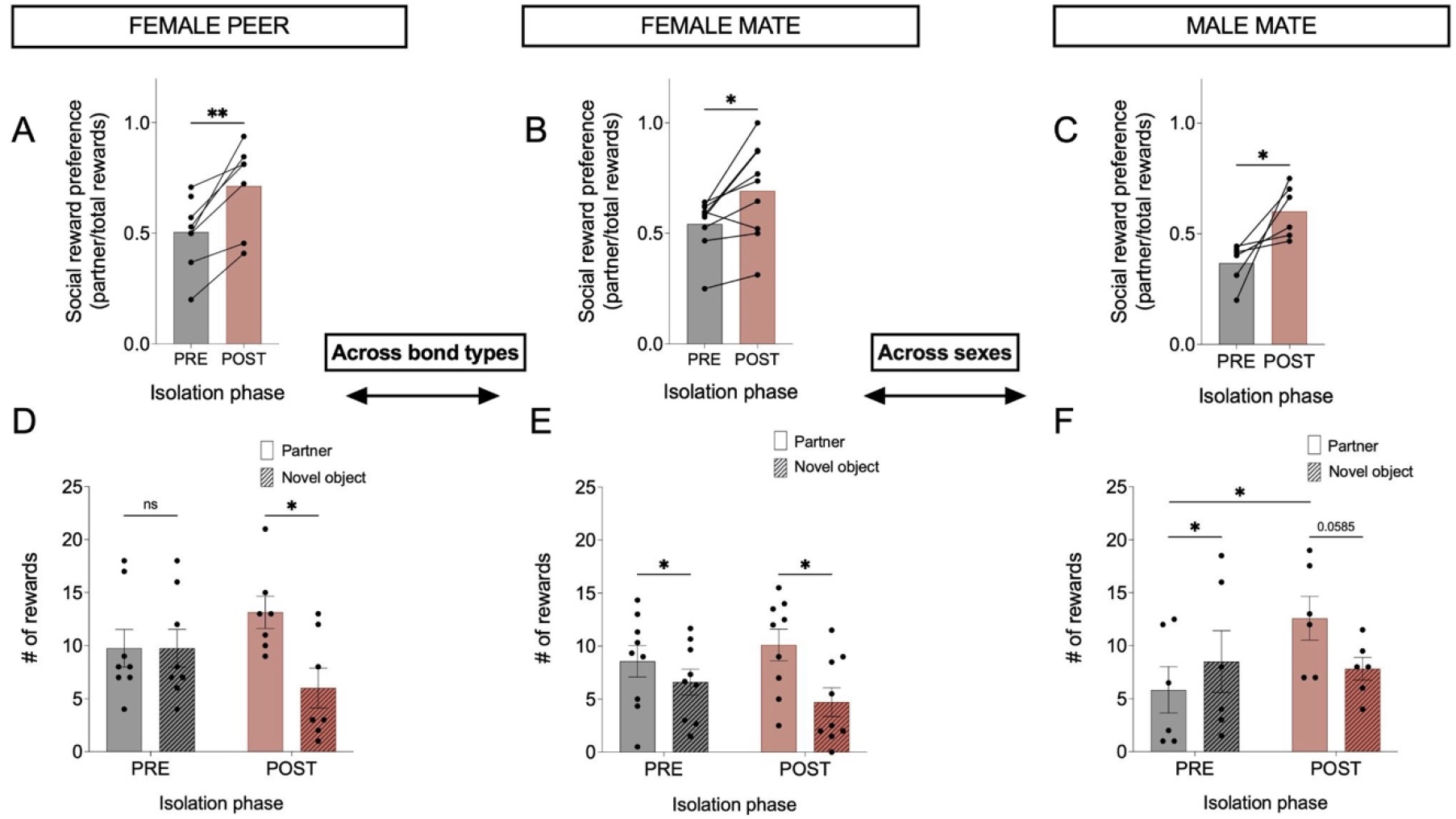
Separation rapidly elevated partner-directed motivation across relationship types and sexes. (*A-C*) Social reward preference scores (*D-F*) and raw total rewards by stimulus target displayed pre- (grey) and post- (red) separation in female peer pairs (n=8) (*A, D)*, female mate pairs (n=9) *(B, E)*, and male mate pairs (n=6) *(C, F)*. Brackets indicate pairwise comparisons, and arrows indicate model comparisons for relationship type (female peer vs. female mate) and sex (female mate vs. male mate). Detailed model results are provided in the text and Table S1. * = <.05, ** = <.01, *** <.001, **** < .0001. Error bars denote the standard error of the mean.

To determine if the type of social bond [peer | mate] or focal vole sex [male | female] modulated this response, we used a linear mixed model (LMM) for each comparative framework. Our model comparing relationship type revealed no difference in social reward preference score between females in peer partnerships and females in mate partnerships (F (1,15)=0.000, *p*=0.9973), displaying only an effect of isolation phase (F(1,14.3)=30.3661, p<.0001). The sex-comparison model highlighted a marginal effect of focal sex (F(1,13)=3.8288, *p*=0.0722) and a strong effect of isolation phase (F(1,13)=18.9744, *p*=0.0008). Together, these analyses indicate that partner separation robustly increased the motivation to access the lost partner in both sexes regardless of whether the partner was a mate or peer.

### Baseline social reward differed by focal sex, but partner separation increased social motivation in all groups

We next evaluated the raw number of rewards using two models accounting for the group comparison (bond type [peer | mate] *or* focal sex [male | female]), isolation condition, stimulus target [partner | novel object], and their interactions (*Figure 1D-F*).

The model across relationship type revealed a significant main effect of reward target (F(1,59)=26.2717, *p*<.0001) and a target*isolation phase interaction (F(1,59)=13.6429, *p*=0.0005), with no differences in number of rewards by bond type. Further pairwise comparisons showed that while female peers (*D*) worked equally hard for both stimuli at baseline (t(7)=0, *p*>0.999), female mates (*E*) worked harder to access their male partner even before isolation (t(8)=2.706, *p*=0.02681). Following partner loss, females of both relationship types worked harder to access the social reward versus the novel object after separation from their partner (peers: t(6)=2.634, *p*=0.03885; mates: t(8)=2.890, *p*=0.02021).

The model evaluating focal sex highlighted a main effect of stimulus target (F(1,53)=9.54992, *p*=0.0032) on raw reward count, alongside significant interactions between isolation phase*target (F(1,53)=11.9934, *p*=0.0011), sex*target (F(1,53)=5.66027, *p*=0.0210), and sex*isolation phase (F(1,53)=6.8294, *p*=0.0116). In contrast to the female cohorts, male voles (*F*) were more motivated by the novel object at baseline (t(5)=3.280, *p*=0.02196). Still, they increased their partner-directed rewards following social loss (t(5)=2.2395, *p*=0.0491), developing a trend towards separation-induced preference for the social reward over the novel object (t(5)=2.443, *p*=0.05847). Thus, despite marked differences in baseline social reward across sex and relationship type, partner loss consistently shifted motivation toward the absent social partner across all groups.

### Separation from a bonded partner elevated stimulated dopamine release

We next sought to investigate whether enhanced motivation for social reward following partner separation correlated with changes in the signaling of dopamine, a neuromodulator involved in motivation. For this purpose, we compared stimulated dopamine release in *ex vivo* brain slices from voles following social loss and their non-isolated control counterparts. To characterize dopamine release and clearance dynamics, we used near-infrared catecholamine nanosensors (nIRCats; *Figure S1A*), which have previously been used to image dopamine signaling with high spatiotemporal resolution in *ex vivo* mouse (59), meadow vole (61), and prairie vole (58) brain slices. We applied nIRCats to brain slices containing the nucleus accumbens (NAc) (*Figure S1B, C*), a region in which altered dopamine signaling has been observed after long-term social isolation in rats, and where dopamine responses to partners and strangers differ using ensemble-level measurements in prairie voles in stable mate partnerships (53, 56). While the NAc receives multiple neurochemical inputs, nIRCats are selective for dopamine and norepinephrine over other molecules (59), and the NAc lacks norepinephrine inputs (62).

Electrical stimulation evoked a robust increase in nIRCat fluorescence (ΔF/F_0_) that returned to baseline (*Figure S1D, E*), allowing us to first characterize the ΔF/F_0_ integrated across the entire field of view (FOV [140 µm × 175 µm]), as a measure of evoked dopamine release. Statistical model analysis was conducted in two stages. We first evaluated the fixed effects of treatment, bond type, and their interaction exclusively within the female dataset because males had only one relationship type. Model comparison revealed that a simplified model excluding bond type provided a superior fit (ΔAICc > 2) for all imaging metrics. Thus, female relationship types were pooled for all subsequent analyses in which we analyzed the combined dataset (males and pooled females) using a linear mixed model with isolation phase, sex, and their interaction, including subject ID to account for multiple samples (separate hemispheres or slices) obtained from the same animal (*Table S2*). Using nIRCat imaging, we found that voles that lost a bonded partner exhibited a higher dopamine release response compared to unseparated controls, a pattern observed in both females and males (*Figure 2A-D*). The model further revealed a significant effect of isolation condition (f(1,76)=5.569, *p*=0.0208) and focal sex (f(1,76)=7.3509, *p*=0.0083) on integrated ΔF/F_0_ upon stimulation (*Figure S2*), indicating that acute partner loss enhances evoked NAc dopamine release across sexes and relationship types following social separation.

**Figure 2.**
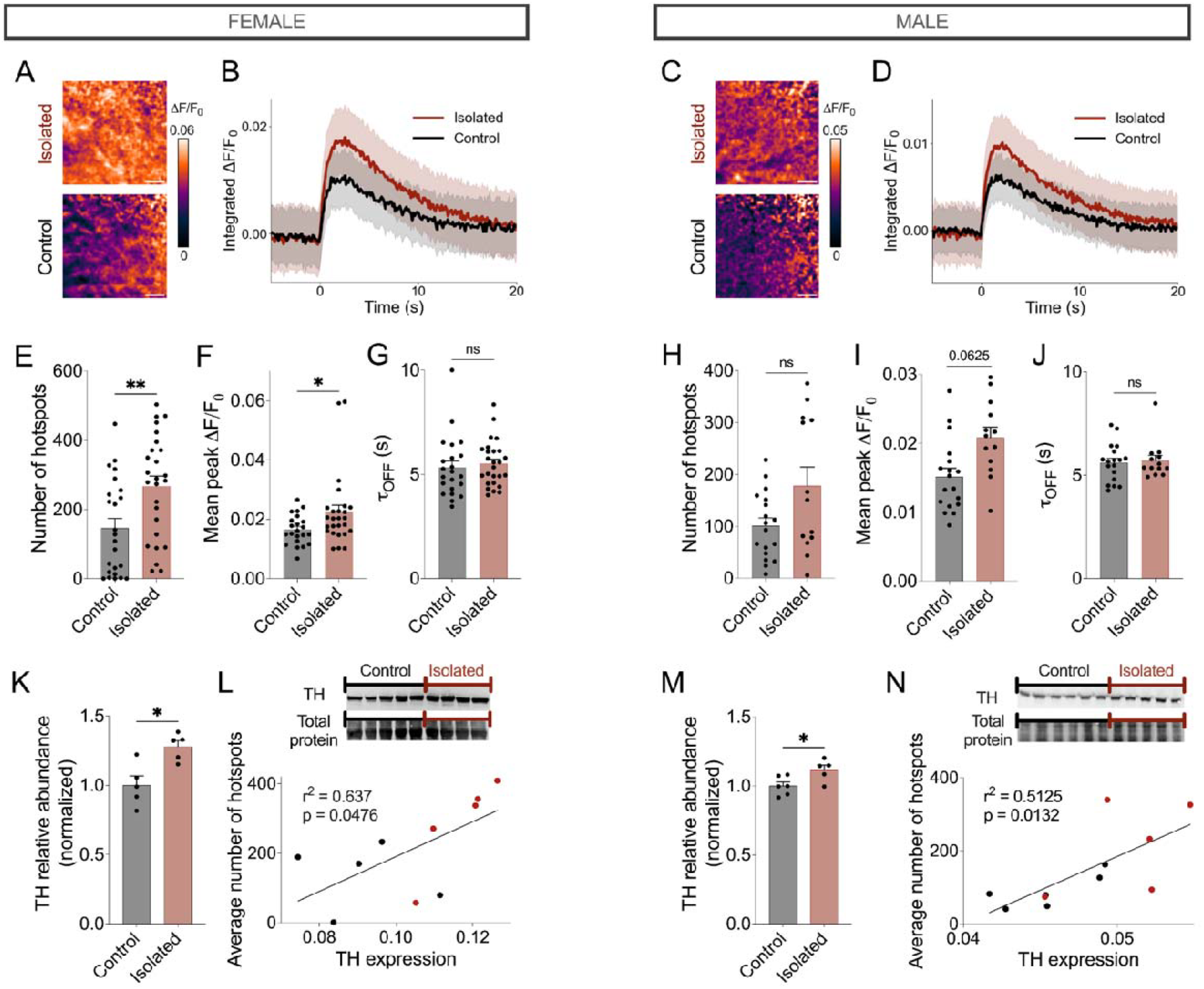
Separation enhanced mesolimbic dopamine signaling in male and female voles. (*A-J*) Evoked dopamine release in the nucleus accumbens imaged with nIRCats. (*A, C*) Representative ΔF/F_0_ images upon stimulation for the isolated (top) and control group (bottom) of female (*A*) and male (*C*) voles. Scale bars represent 10 μm. (*B, D*) Time course of integrated ΔF/F_0_ in isolated (red) and control group (black) of female (*B*) and male (*D*) voles. The solid line denotes the averaged value, and the shading indicates the standard deviation (SD). (*E, H*) Number of hotspots identified in the field of view for female (*E*) and male (*H*) voles. (*F, I*) ΔF/F_0_ averaged over all hotspots for female (*F*) and male (*I*) voles. (*G, J*) ΔF/F_0_ signal decay constant _OFF_ averaged over all hotspots for female (*G*) and male (*J*) voles. (*A-J*) n = 24 samples (11 animals) for control and n = 26 samples (12 animals) for isolated groups in female voles, and n = 18 samples (8 animals) for control and n = 13 samples (6 animals) for isolated groups in male voles. See relationship type comparison within female voles in *Figure S3* and sex comparison within mate relationships in *Figure S4*. (*K-N*) Tyrosine hydroxylase (TH) levels measured by Western blot. (*K, M*) TH protein normalized to the total protein of the respective lane for female (*K*) and male (*M*) voles and displayed as relative abundance to the control mean. See nonnormalized, raw images in *Figure S5*. (*L, N*) Average hotspots per vole against TH level for female (*L*) and male (*N*) subjects in control (black) and isolated (red) groups. Fit (black solid line) by simple linear regression. * = <.05, ** = <.01, *** <.001, **** < .0001 (see *Table S2* for full statistics). Error bars denote the standard error of the mean.

### Subcellular analysis revealed a separation-induced increase in the number of dopamine release hotspots

To resolve subcellular dopamine release dynamics, we leveraged the high spatial resolution of nIRCat (2 µm) by applying a spatial grid-filtering analysis to identify discrete dopamine “hotspots” where peak ΔF/F_0_ following stimulation exceeded three times the standard deviation of the baseline (61) (*Figure S1F*). Each hotspot has been associated with localized dopamine release from an individual varicosity in neuronal culture (63); therefore, the number of hotspots identified serves as a proxy for the number of putative active dopamine release sites per field of view. Within each hotspot, we can calculate additional metrics such as hotspot peak ΔF/F_0_ and τ_OFF_, which, when averaged across all hotspots, can be used to assess the relative amount of dopamine released per hotspot and the local clearance time, respectively. Evaluating release and reuptake dynamics at individual sites offers a highly sensitive metric for detecting changes in neurochemical signaling across experimental cohorts (60).

Separation from a partner resulted in a greater number of evoked dopamine hotspots (*Figure 2E, H*; f(1,76)=9.4391, *p*=0.0029) as well as greater mean peak ΔF/F_0_ within hotspots (*Figure 2F, I*; f(1,74)=8.934, *p*=0.0038) compared to controls. Focal sex significantly influenced the number of hotspots (f(1,76)=6.7964, *p*=0.011), but did not affect the magnitude of release (ΔF/F_0_) across hotspots (f(1,74)=0.9887, *p*=0.7541). There was no statistically significant difference in the mean τ_OFF_ within hotspots (*Figure 2G, J*) by isolation condition (f(1,74)=0.2796, *p*=0.5985) or sex (f(1,74)=0.2871, *p*=0.5937). These results demonstrate that partner separation enhances presynaptic dopamine signaling by increasing both the number of active release sites and the amount of dopamine released per site, without altering local dopamine clearance.

### Separation increased tyrosine hydroxylase abundance, correlating with release site density

To identify the molecular underpinnings of elevated dopamine release following separation, we quantified striatal tyrosine hydroxylase (TH) levels using Western blotting. TH levels were normalized to total protein loading per lane. Females subjected to partner separation exhibited higher normalized TH abundance compared to non-separated controls (*Figure 2K*; f(1,6)=8.1817, *p*=0.0288, two-way ANOVA), with no main effect of relationship type. For males, where pair type was not a factor, a pairwise comparison likewise revealed a significant elevation in TH levels following separation (*Figure 2M*; t(7)=2.550, *p*=0.0343).

Given the separation-induced increase in putative dopamine release sites, we sought to investigate whether the increased number of dopamine release sites was accompanied by structural plasticity. To this end, we quantified β - actin, a cytoskeletal protein involved in neuronal remodeling and terminal growth. We found that the normalized abundance of the cytoskeletal protein β - actin was also elevated by bond loss in females (f(1,6)=158.7203, *p*< .0001) and was higher in separated mate pairs relative to separated peer pairs (f(1,6)=17.6507, *p*=0.0057). In contrast, partner loss did not affect β - actin levels in males (t(7)=1.2468, *p*=0.2511). Finally, striatal TH positively correlated with the average number of hotspots identified via nIRCat imaging in both females (*Figure 2L*; f(1,8)=5.463, *p*=0.0476) and males (*Figure 2N*; f(1,9)=9.460, *p*=0.0132). Together, these findings suggest that the increased density of active dopamine release sites following partner loss is associated with enhanced dopaminergic synthetic capacity and, at least in females, cytoskeletal remodeling consistent with structural plasticity.

## Discussion

The loss or disruption of social relationships is fundamentally distressing and can lead to severe health consequences. In this study, we provide the first characterization of how social separation from bonded partners affects effortful social motivation and dopamine release in male and female prairie voles, and the first direct measurement of dopamine release following loss at the subcellular level. While the effects of social isolation on behavior and dopamine signaling have been extensively studied in mice and rats, these models do not display selective preferences for familiar individuals (18, 64, 65). The socially monogamous prairie vole provides a rare opportunity to study both the effect of social separation from *bonded* partners in both sexes and to compare the effects of bond loss across different types of relationships.

### Baseline partner seeking differed by sex and relationship type

We found that baseline social motivation varied by both sex and relationship type. While female voles worked harder to access a mate versus a novel object, they had no such preference for a peer companion, whereas males expended more effort to access the novel object reward over their mate. This sex difference in partner-directed motivation in stably paired voles mirrors our prior studies of social motivation in which only female prairie voles exhibit selective social reward for a mate or peer partner versus a novel conspecific (20, 57, 66). These findings reinforce the idea that selective social bonds do not necessarily confer equivalent motivational value across sexes or relationship types, even within a species in which both sexes form pair-bonds. Importantly, differences in operant social reward do not imply differences in bond selectivity, as male prairie voles exhibit robust partner preferences for both peers and mates despite relatively weak partner-directed operant motivation (20, 57, 66).

### Partner loss ubiquitously enhanced social motivation across sex and relationship type

We used a lever-pressing choice paradigm to directly quantify social reward before and after partner separation in prairie voles. Unlike measures of partner preference or investigation, this approach quantifies the effort an animal is willing to expend to regain access to its partner, providing a direct measure of the motivational value of social reunion. Despite baseline differences in relative pressing for the partner across groups, all prairie voles worked harder to access their bonded partner after five days of separation compared to pre-separation. The observed increase in social motivation indicates a strong drive for reunion following partner loss. However, the enhanced motivation for partner reunion cannot be decoupled from the general drive for social interaction following isolation, as both interpretations align with the social homeostasis hypothesis (67), where deprivation of essential needs (in this case, social interaction) leads to goal-directed behaviors to restore homeostasis. While a recent study demonstrated that brief (3-day) isolation in adult prairie voles does not produce increased affiliative contact with an unfamiliar same- or opposite-sex stranger during free social interactions (51), the increased appetitive social drive we see for for reunion with the separated partner may result from use of a bonded partner as the social stimulus, from the longer duration of separation, or from a different testing paradigm. This aligns with prior work showing that post-separation increases in homecage bedding investigation do not generalize to unfamiliar animal odor cues (31, 50).

Furthermore, the uniform increase in social seeking behavior across both reproductive and non-reproductive relationships highlights the fundamental importance of both bond types for prairie voles. This finding underscores the salience of female peer relationships, as their loss evoked a neurobehavioral response similar to the disruption of a reproductive pair bond. Additionally, the increase in mate-directed seeking across both males and females indicates the importance of bonded mate relationships across sexes, consistent with the species’ socially monogamous mating system.

### Partner loss elevated dopamine release

Given the central role of mesolimbic dopamine in social reward and motivated behavior, we sought to determine how partner loss alters dopamine release itself and, using the high spatial resolution of nIRCat nanosensors, whether these adaptations occur at the level of individual presynaptic release sites. We observed elevated dopamine release in the NAc following social separation in all contexts. This finding aligns with prior work in rodent models, including an *ex vivo* fast-scan cyclic voltammetry (FSCV) study showing increased dopamine release in the NAc core after six weeks of isolation in rats (53), and *in vivo* microdialysis evidence reporting heightened accumbal dopamine release following early-life isolation in mandarin voles (54). However, these studies either focused exclusively on males in non-monogamous species or relied on developmental and long-term (> 2-week) isolation paradigms. By evaluating both male and female prairie voles across distinct bond types following 5 days of separation in adulthood, our study demonstrates that elevated NAc dopamine signaling is a rapid and widespread signature of social loss across diverse contexts.

The high spatial resolution (∼2 µm) of nIRCats is comparable to the size of varicosities in dopaminergic neurons (59), enabling the first quantification of separation-induced effects on dopamine release hotspot density. These hotspots can be used to approximate the number of dopamine release sites, and the peak ΔF/F_0_ in each hotspot can represent the amount of dopamine released per site (58, 61), both of which have been validated in dopaminergic neuron culture studies (63). In our dopamine hotspot analysis, we observed a larger number of release sites in separated groups relative to control groups, suggesting a possibility of structural changes in neuron morphology or physiology induced by acute partner loss. We further observed an increase in the mean peak ΔF/F_0_ from identified dopamine hotspots in both sexes following social loss, suggesting that varicosities in the separated group also release more dopamine per site. Finally, we found that dopamine clearance remained unaffected by isolation conditions, suggesting that partner separation may predominantly affect dopamine synthesis and release rather than reuptake kinetics moderated by dopamine transporters.

These findings align with previous work in voles demonstrating that both acute and chronic social isolation alter postsynaptic markers of dopamine signaling in the NAc. Specifically, isolation increases dopamine receptor D1 mRNA (50, 54) and protein expression (68) in both sexes while decreasing dopamine receptor D2 mRNA in females (54). While previous studies primarily focused on postsynaptic targets such as dopamine receptors, the use of nIRCat nanosensors enabled direct monitoring of evoked dopamine release itself. To further substantiate our presynaptic observations of increased dopamine release-site density and increased dopamine release per release site, we quantified TH abundance in striatal tissue by Western blot analysis, which revealed an increase in the rate-limiting enzyme for dopamine synthesis following partner separation. Moreover, TH abundance is positively correlated with dopamine release-site density, suggesting that enhanced dopamine synthetic capacity may support the recruitment or maintenance of additional active release sites following social loss. Together with the above evidence for postsynaptic dopamine receptor adaptations in the NAc, these findings suggest that social loss also engages presynaptic dopaminergic plasticity, revealing coordinated adaptations across both sides of dopamine signaling.

### Separation induced a local increase in dopamine synthesis

To further understand the separation-induced increase in dopamine release and the increase in active release site density, we characterized the abundance of tyrosine hydroxylase (TH) and β - actin in the NAc and dorsal striatum using Western blotting. Because these brain regions lack norepinephrine inputs (62), TH can be used as a marker of dopaminergic terminals and dopamine synthetic capacity. We observed increased levels of TH following separation in both males and females, which aligns with prior work showing heightened accumbal TH levels in isolated male rats (69). Elevated TH levels provide evidence that increases in evoked dopamine release are mediated by upregulated dopamine synthesis. Notably, TH abundance positively correlated with dopamine release-site density within the same animals, linking increased dopamine synthetic capacity to the greater number of active release sites observed following partner separation in both sexes. These data suggest that the greater density of active dopamine release sites following social separation may stem from increased local dopamine synthesis, recruitment of previously inactive release sites, or structural remodeling of dopaminergic terminals–consistent with prior evidence that acute social isolation elevates dopamine neuron excitability (55).

Total protein-normalized Western blots also revealed an upregulation in β - actin in voles that lost their bonded partners. While β - actin is traditionally utilized as a loading control, treatment-driven shifts in the cytoskeletal protein are associated with neural plasticity and changes in neuronal structure such as terminal growth (70). Therefore, the separation-induced elevation of β - actin may align with the increased dopamine release site density following bond loss. Future studies examining presynaptic release machinery, including Bassoon, RIM, and Munc13, could distinguish whether the increased number of active release sites reflects functional recruitment of existing terminals or structural remodeling following social loss (71).

In summary, separation from a bonded partner—whether a mate partner or peer partner— induces a profound shift in the behavioral and neurochemical measures of social reward in both male and female prairie voles. By integrating an operant social choice paradigm with high-resolution dopamine imaging, we demonstrate that partner loss enhances social motivation, mirrored by increases in evoked dopamine release and the proliferation of active release sites within the NAc. Increased TH abundance and its correlation with release-site density further suggest that enhanced dopamine synthetic capacity accompanies these functional adaptations. These results suggest that—across relationship types and sexes—the brain compensates for loss of a bonded partner by amplifying mesolimbic reward system sensitivity and output.

## Materials and Methods

### Animal subjects

Prairie voles were bred in a long photoperiod (14 hr light:10 hr dark). Voles were group-weaned at postnatal day (PND) 21 and separated into same-sex pairs by PND 28. In adulthood, subjects were paired with a new same-sex peer partner or opposite-sex mate partner for 12-16 days prior to the start of behavioral testing or imaging. All experimental procedures were conducted according to federal and institutional guidelines, and protocols were approved by the Animal Care and Use Committee of UC Berkeley.

### Experiment 1. Operant-conditioned lever pressing

Voles were tested in an operant social choice assay to assess changes in social motivation following loss of a bonded partner. Testing consisted of 3 groups: female voles with their peer partner (n=8), female voles with a mate partner (n=9), and male voles with their mate partner (n=6). Training began between PND 45-55 and was conducted according to a previously described protocol (57, 58). Following training, voles began social choice testing with the vole’s partner tethered in one chamber and a novel object in a second chamber. Social choice testing consisted of two phases: (1) pre-separation: partner versus novel object and (2) post-separation: partner versus novel object. Prior to separation, focal voles were exposed to operant social choice for a minimum of 5 days, or until performance stabilized. The order of novel objects was balanced across separation phases and remained consistent across experimental groups. During each 30-minute trial, the vole could access a specific reward by lever-pressing four times on that side, opening the door to the respective stimulus chamber. After one minute, the door closed, and the experimenter moved the focal animal back to the center chamber. Opposite sex partners of focal voles were sterilized via tubal ligation or castration with testosterone replacement (as in (20)) 10-15 days before placement in mate pairs with intact focal animals.

The number of lever presses and rewards during the testing session was recorded with MED-PC VI software (Med Associates). Social reward preference scores were calculated as the number of social rewards out of the total rewards (social plus novel object). Testing sessions were included only if the animal pressed each lever at least once. Across all experiments, a total of five trials did not meet this criterion and the adjacent day of testing was used instead.

### Experiment 2. *Ex vivo* dopamine imaging

In order to assess the effects of relationship loss in different relationship types and across the sexes, we conducted *ex vivo* imaging of dopamine release in control (non-separated) and separated conditions across 3 groups: female peers (n=22 samples from 11 animals), female mates (n=28 samples from 12 animals), and male mates (n=31 samples from 14 animals).

#### Synthesis of nanosensors and brain slice labeling

nIRCat nanosensors were prepared as described previously (59). Briefly, HiPCo SWCNT slurry (NanoIntegris) and ssDNA ((GT)_6_, Integrated DNA Technologies) were mixed at a 1:2 mass ratio in 10 mM NaCl water and probe-tip sonicated (Cole-Parmer Ultrasonic Processor, 3-mm tip) for 30min at 50% amplitude. The resulting suspensions were centrifuged at 21,000 g for 240 min to pellet unsuspended SWCNT.

Acute brain slices were prepared following the protocols described previously (59). Briefly, after transcardial perfusion, the extracted brain was cut into 300 µm thick coronal slices (Leica VT 1000). Three slices per animal containing the nucleus accumbens were prepared. Two hemispheres from the central slice were used for imaging, and the first and last slices were used for Western blot protein extraction. Slices were incubated at 37°C for 30 min in carbogen (95% O_2_, 5% CO_2_, Praxair)-saturated aCSF (in mM as follows: 119 NaCl, 2.5 KCl, 1.3 MgCl_2_, 1 NaH_2_PO_4_, 26.2 NaHCO_3_, 10 glucose, and 2 CaCl_2_) in a chamber (Scientific Systems Design, Inc., BSK4) and then transferred to room temperature for 30 min. Nanosensors were applied to the surface of brain slices with a pipette to a final concentration of 2 mg/L and incubated for 15 min.

#### nIR microscope design and stimulation-evoked imaging

A custom upright epifluorescence microscope (Olympus, Sutter Instruments) described in greater detail previously (59, 60) was used to image nanosensor fluorescence response. Briefly, a 785 nm excitation laser (OptoEngine LLC, MDL-III-785R-300mW) was used to excite nanosensors, and nanosensor fluorescence was collected with a two-dimensional InGaAs array detector (Raptor Photonics, Ninox 640). Microscope operation was controlled by Micro-Manager Open Source Microscopy Software 42. Carbogen-bubbled aCSF was flowed through the microscope at a rate of 2 mL/min. Imaging chamber temperature was kept at 32°C (Warner Instruments, TV-324C). Slices were placed in the chamber with a tissue harp, and a bipolar stimulation electrode (MicroProbes for Life Science, PI2ST30.1A5) was positioned at least 80 µm away from the target field of view. nIR fluorescence images were acquired at frame rates of 8 frames/s (nominal) for 600 frames, where 1 millisecond of 0.1 mA stimulation was applied at the 200th frame. This image acquisition was repeated three times, with 5 min of waiting time in between.

#### Image processing and data analysis

Files were processed using custom Python code (https://github.com/NicholasOuassil/NanoImgPro). Integrated ΔF/F_0_ (ΔF/F_0_ for the entire field of view, 175 µm by 140 µm) was calculated as ΔF/F_0_ = (F-F_0_) /F_0_, where F_0_ is the average intensity for the first 5% of frames, and F is the dynamic fluorescence intensity from the entire field of view. Next, a 25×25 pixel (corresponding to 6.8 µm by 6.8 µm) grid mask was applied to the image stack, and then a median filter convolution within each grid was calculated (*Figure S1F*). For each grid square, ΔF/F_0_ was calculated, and hotspots were identified if the F at the time of stimulation (200 frames) was 3 standard deviations above the baseline F_0_ activity.

### Experiment 3. Western blot protein quantification

To assess the impact of bond loss on proteins associated with dopamine signaling and cytoskeletal remodeling, we quantified tyrosine hydroxylase and β - actin protein levels relative to total protein in male and female brain slices following 5 days of separation (n=6 male, 5 female) and compared them to unseparated controls (n=5 male, 5 female). For each vole, the dorsal and ventral striatum were microdissected bilaterally from the two 300 μL slices directly adjacent to the slice used for nIRCat sensor imaging, then immediately flash-frozen. Tissue samples were homogenized and sonicated with 300 μL of RIPA buffer and 9 μL of 3% Halt protease and phosphatase inhibitor cocktail. Protein quantification was determined at a 2x dilution using the Pierce Rapid Gold Protein Assay Kit (Pierce Biotechnology) according to the corresponding protocol. Samples were mixed with lithium dodecyl sulfate buffer containing β-mercaptoethanol and denatured at 70°C for 10 min before storage at −80 °C. 20ug of protein and 5 µL of PageRuler Prestained NIR Ladder were loaded into a 4-12% gradient tris-bis Bolt gel prior to transfer to nitrocellulose membranes. Protein transfer was confirmed using Revert total protein stain. Membranes were blocked for 1 hour at room temperature (or overnight at 4 degrees) in Intercept Blocking Buffer (LICORbio, 92760001). Membranes were then incubated at 4 degrees overnight with antibodies against tyrosine hydroxylase (mouse anti-TH, MAB318MI, 1:1000). Finally, membranes were incubated with IRDye 800CW secondary antibodies (goat anti-mouse, LICORbio #926-32280, 1:10,000) and visualized using an Azure 500 imaging system (Azure Biosystems). After imaging, the membranes were stripped, re-probed, and visualized using 1x Reblot Plus (2504), antibodies against β-actin (mouse anti-actin, A1978, 1:10,000), and IRDye 800CW secondary antibodies (goat anti-mouse, LICORbio #926-32280, 1:20,000). Protein band intensities were quantified in ImageJ 1.54 and normalized to the total protein abundance per lane.

### Statistical analysis

Social reward preference scores were analyzed using Linear Mixed Models (LMM) with Restricted Maximum Likelihood (REML) estimation and Subject ID as a random effect to account for repeated measures. To analyze the raw reward count data, we implemented a Generalized Linear Mixed Model (GLMM) with a Poisson distribution and Subject ID as a random effect. Fixed effects for the models included isolation phase [pre | post separation], stimulus target [partner | novel object], and group (relationship type [peer | mate] or sex [male | female]). Pairwise comparisons were performed using paired *t*-tests for matched within-subject data and Welch’s *t*- tests for unmatched between-subject groups. nIRCat imaging data were analyzed using an LMM including isolation phase, sex, and their interaction, as well as Subject ID. Model selection was guided by the corrected Akaike Information Criterion (AICc). All statistical modeling and parsing were performed using JMP (version 18) and GraphPad Prism (version 11).

## Acknowledgments

We appreciate Jon Bunting, Daniel Chang Kuo, Lukas Hammett, and Eric Xu for assistance with operant food training and testing, as well as Elizabeth Piotrowski for her contributions to training in protein extraction, quantification, and analysis. We also thank UC Berkeley OLAC for routine animal care. We acknowledge the support of a National Institutes of Health grant R01MH132908 (A.K.B.), a National Science Foundation CAREER award 2239635 (A.K.B.), a McKnight Foundation award (M.P.L.), a Simons Foundation award (M.P.L.), a Moore Foundation award (M.P.L.), a Heising-Simons Fellowship (M.P.L.), a Brain Foundation award (M.P.L.), a Polymaths award from Schmidt Sciences, LLC (M.P.L.), an NSF Biophotonics award (M.P.L), the Rennie Fund (M.P.L), Schmidt Science Fellows, in partnership with the Rhodes Trust (N.K.), a Burroughs Wellcome Fund Career Award at the Scientific Interface (N.K.), and a National Institutes of Health fellowship F31MH141997 (A.M.B). M.P.L. is a Chan Zuckerberg Biohub investigator.

