## Supplementary material for "Acute partner loss enhances social motivation and nucleus accumbens dopamine release in prairie voles": Figure S1A

**This PDF file includes:**

Tables S1 to S4

Figures S1 to S5

Tables

| Comparison | Model & Terms | Test stat | P value |
| --- | --- | --- | --- |
| Pair type comparison of reward ratios (Fig. 1A-B) | **LMM** -2LL=-13.51  Pair type  **Isolation phase**  Pair*isolation  Subject | F(1,15)=0.00  **F(1,14.3)=30.36**  F(1,14.3)=1.23 | 0.9973  **<.0001^****^**  0.2862 |
| Sex comparison of reward ratios (Fig. 1B-C) | **LMM** -2LL=-12.41  Sex  **Isolation phase**  Sex*isolation  Subject | F(1,13)=3.83  **F(1,13)=18.97**  F(1,13)=0.92 | 0.0722  **0.0008^***^**  0.3548 |
| Pair type comparison of total rewards (Fig. 1D-E) | **GLMM (Poisson)** X^2^/DF=1.42  Pair type  Isolation phase  **Target**  Pair*isolation  Pair*target  **Isolation*target**  Subject | F(1,14.7)=1.74  F(1,59)=2.26  **F(1,59)=26.27**  F(1,59)=0.040  F(1,59)=0.60  F(1,59)=13.64 | 0.2178  0.1379  **<.0001^****^**  0.8422  0.4416  0.0005^***^ |
| Sex comparison of total rewards (Fig. E-F) | **GLMM (Poisson)** X^2^/DF=1.324  Sex  Isolation phase  **Target**  **Sex*isolation**  **Sex*target**  **Isolation*target**  Subject | F(1,12.6)=0.25  F(1,53)=1.03  **F(1,53)=9.55**  **F(1,53)=6.83**  **F(1,53)=5.66**  F(1,53)=11.99 | 0.6240  0.3142  **0.0032^**^**  **0.0116^*^**  **0.0210^*^**  0.0011^**^ |

Table S1. Model outputs for operant behavioral data. This table shows all model outputs relating to *Figure 1*. Rows 1- 2 show outputs for social reward preference scores (*Figure 1A-C*). Rows 2-4 show outputs for raw number of rewards by target [partner | novel object] (*Figure 1D-F*). *  =  <.05, **  = <.01, *** <.001, **** < .0001.

| Comparison | Model & Terms | Test stat | P value |
| --- | --- | --- | --- |
| Integrated ΔF/F_0_ (Fig. 2B, D) | **LMM**  AICc -529  **Isolation**  **Sex**  Isolation*Sex  Subject  ***Post hoc* *t*-tests**  **Female (control vs isolated)**  Male (control vs isolated)  **Control (male vs female)**  Isolated (male vs female) | **F(1,76)=5.5695**  **F(1,76)=7.3509**  F(1,76)=0.7383  F(1,76)=0.9929  **t(76)=2.63**  t(76)=0.97  **t(76)=1.99**  t(76)=2.71 | **0.0208^*^**  **0.0083^**^**  0.3929  0.3222  **0.0103^*^**  0.3363  **0.0500^*^**  0.0083^**^ |
| Number of hotspots (Fig. 2E, H) | **LMM** AICc 1021  **Isolation**  **Sex**  Isolation*Sex  Subject  ***Post hoc t*-tests**  **Female (control vs isolated)**  Male (control vs isolated)  Control (male vs female)  Isolated (male vs female) | **F(1,76)=9.4391**  **F(1,76)=6.7964**  F(1,76)=0.6264  F(1,76)=2.4926  **t(76)=3.16**  t(76)=1.47  t(76)=1.93  t(76)=2.59 | **0.0029^**^**  **0.0110^*^**  0.4311  0.1185  **0.0023^**^**  0.1470  0.0570  0.0115 |
| Mean peak ΔF/F_0_ (Fig. 2F, I) | **LMM** AICc -525  **Isolation**  Sex  Isolation*Sex  Subject  ***Post hoc t*-tests**  **Female (control vs isolated)**  Male (control vs isolated)  Control (male vs female)  Isolated (male vs female) | **F(1,74)=8.9340**  F(1,74)=0.0988  F(1,74)=0.0027  F(1,74)=0.0236  **t(74)=2.46**  t(74)=1.89  t(74)=0.25  t(74)=0.29 | **0.0038^**^**  0.7541  0.9582  0.8782  **0.0162^*^**  0.0625  0.8006  0.7706 |
| $\boldsymbol{\tau}_{\mathbf{OFF}}$ (Fig. 2G, J) | **LMM** AICc 256  Isolation  Sex  Isolation*Sex  Subject  ***Post hoc t*-tests**  Female (control vs isolated)  Male (control vs isolated)  Control (male vs female)  Isolated (male vs female) | F(1,74)=0.2796  F(1,74)=0.2870  F(1,74)=0.0221  F(1,74)=0.0035  t(74)=0.55  t(74)=0.25  t(74)=0.55  t(74)=0.38 | 0.5985  0.5937  0.8822  0.9525  0.5864  0.8062  0.5823  0.7024 |
| Female TH level (Fig. 2K, L) | **2-way ANOVA**  **Isolation**  Pair  Isolation*Pair  **Linear regression** (R^2^=0.41)  TH level by hotspot | **F(1,6)=8.1817**  F(1,6)=0.0392  F(1,6)=0.1693  F(1,8)=5.463 | **0.0288^*^**  0.8496  0.6950  0.0476^*^ |
| Male TH level (Fig. 2M, N) | **Welch’s *t*-test**  **Isolation**  **Linear regression** (R^2^=0.51)  TH level by hotspot | **t(8)=2.550**  F(1,9)=9.460 | **0.0343^*^**  0.0132^*^ |

Table S2. Model outputs for nIRcat dopamine imaging and Western blot analysis. This table shows all model outputs relating to *Figure 2*. *  =  <.05, **  = <.01, *** <.001, **** < .0001.

| Comparison | Model & Terms | Test stat | P value |
| --- | --- | --- | --- |
| Integrated ΔF/F_0_ (Fig. S3A) | **2-way ANOVA**  **Isolation**  Pair type  Isolation*Pair type  ***Post hoc* Fisher’s LSD**  **Peer (control vs isolated)**  Mate (control vs isolated)  Control (peer vs mate)  Isolated (peer vs mate) | **F(1, 46) = 5.976**  F(1, 46) = 2.617  F(1, 46) = 1.047  **t(46)=2.313**  t(46)=1.074  t(46)=0.4101  t(46)=1.917 | **0.0184^*^**  0.1126  0.3117  **0.0253^*^**  0.2884  0.6836  0.0615 |
| Number of hotspots (Fig. S3B) | **2-way ANOVA**  **Isolation**  Pair type  Isolation*Pair type  ***Post hoc* Fisher’s LSD**  **Peer (control vs isolated)**  Mate (control vs isolated)  Control (peer vs mate)  Isolated (peer vs mate) | **F(1,46) = 8.989**  F(1,46) = 0.06428  F(1,46) = 0.4015  **t(46)=2.422**  t(46)=1.787  t(46)=0.2622  t(46)=0.6440 | **0.0044^**^**  0.8010  0.5295  **0.0194^*^**  0.0806  0.7944  0.5228 |
| Mean peak ΔF/F_0_ (Fig. S3C) | **2-way ANOVA**  **Isolation**  Pair type  Isolation*Pair type  ***Post hoc* Fisher’s LSD**  Peer (control vs isolated)  Mate (control vs isolated)  Control (peer vs mate)  Isolated (peer vs mate) | **F(1,44) = 4.298**  F(1,44) = 1.435  F(1,44) = 0.1303  t(44)=1.616  t(44)=1.301  t(44)=0.5649  t(44)=1.160 | **0.0441^*^**  0.2374  0.7198  0.1133  0.1999  0.5750  0.2523 |
| $\boldsymbol{\tau}_{\mathbf{OFF}}$ (Fig. S3D) | **2-way ANOVA**  Isolation  Pair type  Isolation*Pair type  ***Post hoc* Fisher’s LSD**  Peer (control vs isolated)  Mate (control vs isolated)  Control (peer vs mate)  Isolated (peer vs mate) | F(1,44) = 0.1248  F(1,44) = 0.01592  F(1,44) = 0.8821  t(44)=0.3890  t(44)=0.9823  t(44)=0.7193  t(44)=0.6050 | 0.7256  0.9002  0.3527  0.6991  0.3313  0.4758  0.5483 |

Table S3. Model outputs for nIRcat dopamine imaging and for relationship type comparison in female voles. This table shows all model outputs relating to *Figure S3*. *  =  <.05, **  = <.01, *** <.001, **** < .0001

| Comparison | Model & Terms | Test stat | P value |
| --- | --- | --- | --- |
| Integrated ΔF/F_0_ (Fig. S4A) | **2-way ANOVA**  **Isolation**  **Sex**  Isolation*Sex  ***Post hoc* Fisher’s LSD**  Female (control vs isolated)  Male (control vs isolated)  **Control (female vs male)**  Isolated (female vs male) | **F(1,55) = 6.703**  **F(1,55) = 9.930**  F(1,55) = 0.04128  t(55)=1.938  t(55)=1.720  **t(55)=2.170**  t(55)=2.285 | **0.0123^*^**  **0.0026^**^**  0.8397  0.0577  0.0911  **0.0343^*^**  0.0262^*^ |
| Number of hotspots (Fig. S4B) | **2-way ANOVA**  **Isolation**  Sex  Isolation*Sex  ***Post hoc* Fisher’s LSD**  **Female (control vs isolated)**  Male (control vs isolated)  Control (female vs male)  Isolated (female vs male) | **F(1,55) = 7.609**  F(1,55) = 3.820  F(1,55) = 0.1096  **t(55)=2.145**  t(55)=1.750  t(55)=1.195  t(55)=1.557 | **0.0079^**^**  0.0557  0.1627  **0.0364^*^**  0.0857  0.2371  0.1253 |
| Mean peak ΔF/F_0_ (Fig. S4C) | **2-way ANOVA**  **Isolation**  Sex  Isolation*Sex  ***Post hoc* Fisher’s LSD**  **Female (control vs isolated)**  **Male (control vs isolated)**  Control (female vs male)  Isolated (female vs male) | **F(1,54) = 14.13**  F(1,54) = 4.797e-006  F(1,54) = 0.09226  **t(54)=2.377**  **t(54)=2.957**  t(54)=0.2195  t(54)=0.2105 | **0.0004^***^**  0.9983  0.7625  **0.0210^*^**  **0.0046^**^**  0.8271  0.8341 |
| $\boldsymbol{\tau}_{\mathbf{OFF}}$ (Fig. S4D) | **2-way ANOVA**  Isolation  Sex  Isolation*Sex  ***Post hoc* Fisher’s LSD**  Female (control vs isolated)  Male (control vs isolated)  Control (female vs male)  Isolated (female vs male) | F(1,54) = 0.8693  F(1,54) = 0.5853  F(1,54) = 0.3748  t(54)=1.063  t(54)=0.2331  t(54)=1.003  t(54)=0.1052 | 0.3553  0.4476  0.5430  0.2927  0.8166  0.3205  0.9166 |

Table S4. Model outputs for nIRcat dopamine imaging and for sex comparison (male relationship type only). This table shows all model outputs relating to *Figure S4*. *  =  <.05, **  = <.01, *** <.001, **** < .0001

Figures

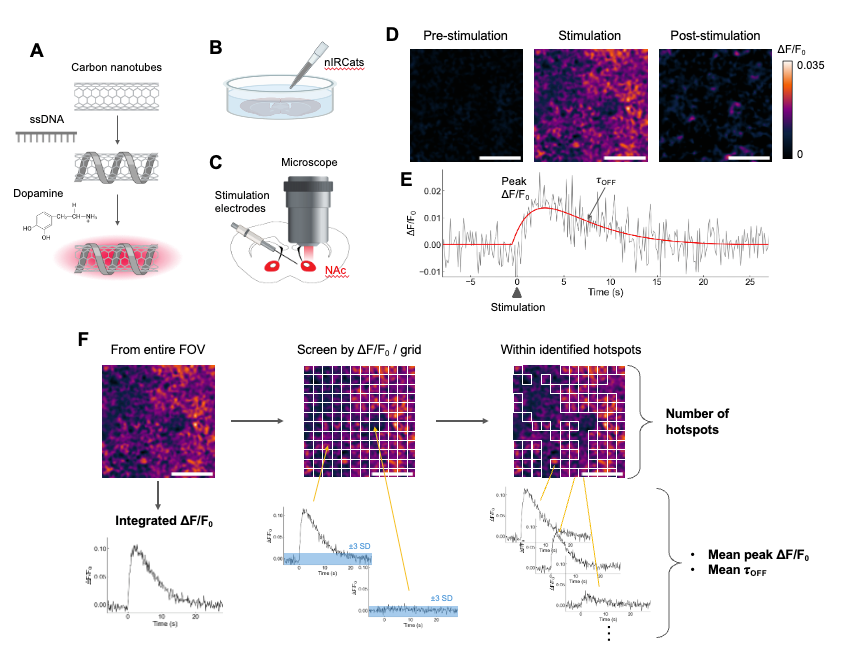

Fig. S1. Dopamine imaging with nIRCat sensors. (*A*) Mechanism of nIRCat. nIRCat is based on single-walled carbon nanotubes functionalized by single-stranded DNA (ssDNA) that confers selectivity for dopamine. A unique ssDNA sequence was identified to modulate the fluorescence intensity of single-walled carbon nanotubes in the presence of dopamine. (*B*) nIRCats in 10 mM NaCl solution were applied by pipettes to acute brain slices. (*C*) We imaged areas within the nucleus accumbens (NAc) and applied electrical stimulation (1 mA) through microelectrodes. (*D*) Representative ΔF/F_0_ images in NAc following 0.1 mA electrical stimulation. Three frames are shown: (left) “pre-stimulation” is the baseline ΔF/F_0_ before electrical stimulation; (center) “stimulation” is immediately after electrical stimulation; and (right) “post-stimulation” is after ΔF/F_0_ has returned to baseline. Scale bars represent 20 μm. (*E*)  Representative time-course ΔF/F_0_ (black solid line). The peak ΔF/F_0_ is identified by first finding the maximum value within 6 seconds (50 frames) from the stimulation, and taking the average of the next 5 frames. We fit the time-course ΔF/F_0_ by $\boldsymbol{\alpha\times}\left( \boldsymbol{1-}\mathbf{exp}\left( \boldsymbol{-}\frac{\boldsymbol{x}}{\boldsymbol{\tau}_{\mathbf{ON}}} \right)\boldsymbol{\times}\left( \mathbf{exp}\left( \boldsymbol{-}\frac{\boldsymbol{x}}{\boldsymbol{\tau}_{\mathbf{OFF}}} \right) \right) \right)\boldsymbol{+\beta}$, where is $\boldsymbol{\alpha}$ scale factor, $\boldsymbol{\beta}$ is a constant, $\boldsymbol{\tau}_{\mathbf{ON}}$ is the rise time constant, and $\boldsymbol{\tau}_{\mathbf{OFF}}$ is the decay time constant. $\boldsymbol{\alpha}$, $\boldsymbol{\beta}$, $\boldsymbol{\tau}_{\mathbf{ON}}$, and $\boldsymbol{\tau}_{\mathbf{OFF}}$ were fitting parameters. (*F*) Image processing flow. Imaging movie files to be processed showing the entire field of view (175 μm by 140 μm). Integrated ΔF/F_0_ (ΔF/F_0_ for the entire field of view) was calculated as ΔF/F_0_ = (F- F_0_) /F_0_, where F_0_ is the average intensity for the first 5% of frames and F is the dynamic fluorescence intensity. Next, a 25x25 pixel (corresponding to 6.8 μm by 6.8 μm) grid mask was applied to the image stack. For each grid square, ΔF/F_0_ was calculated. Grid squares were identified as “hotspot” if the F around time of stimulation (200 frames) is 3 standard deviations above the baseline F_0_ activity. Subsequently, ΔF/F_0_ and $\boldsymbol{\tau}_{\mathbf{OFF}}$ were averaged over all identified hotspots, leading to mean peak ΔF/F_0_ and mean $\boldsymbol{\tau}_{\mathbf{OFF}}$.

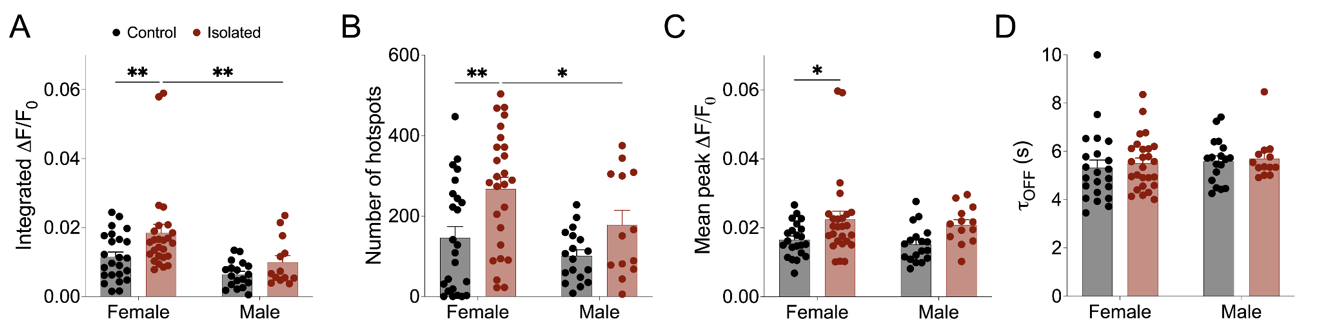

Fig. S2. Sex comparison in male and pooled female voles for parameters from nIRCat imaging. (*A*) Integrated ΔF/F_0_ in isolated (red) and control group (black) of female voles. (*B*) The number of hotspots identified in the field of view. (*C*) ΔF/F_0_ averaged over identified hotspots. (*D*) ΔF/F_0_ signal decay constant _OFF_ averaged over identified hotspots. n = 24 slices (11 animals) for control and n = 26 slices (12 animals) for female, and n = 18 slices (8 animals) for control and n = 13 slices (6 animals) for isolated groups for male. Non-significant comparisons are omitted for clarity. *  =  <.05, **  = <.01, *** <.001, **** < .0001 as determined by post-hoc test. See Table S2 for details. Error bars denote the standard error of the mean.

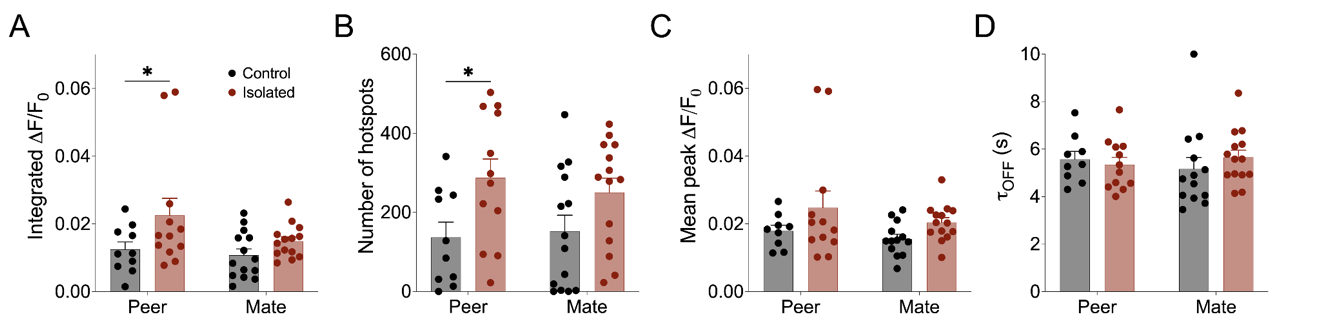

Fig. S3. Relationship type comparison in female voles for parameters from nIRCat imaging. (*A*) Integrated ΔF/F_0_ in isolated (red) and control group (black) of female voles. (*B*) The number of hotspots identified in the field of view. (*C*) ΔF/F_0_ averaged over identified hotspots. (*D*) ΔF/F_0_ signal decay constant _OFF_ averaged over identified hotspots. n = 10 slices (5 animals) for control and n = 12 slices (6 animals) for peer pairs, and n = 14 slices (6 animals) for control and n = 14 slices (6 animals) for isolated groups for mate pairs. Non-significant comparisons are omitted for clarity. *  =  <.05, **  = <.01, *** <.001, **** < .0001 as determined by post-hoc Fisher’s LSD test. See *Table S3* for details. Error bars denote the standard error of the mean. See Table S3 for details.

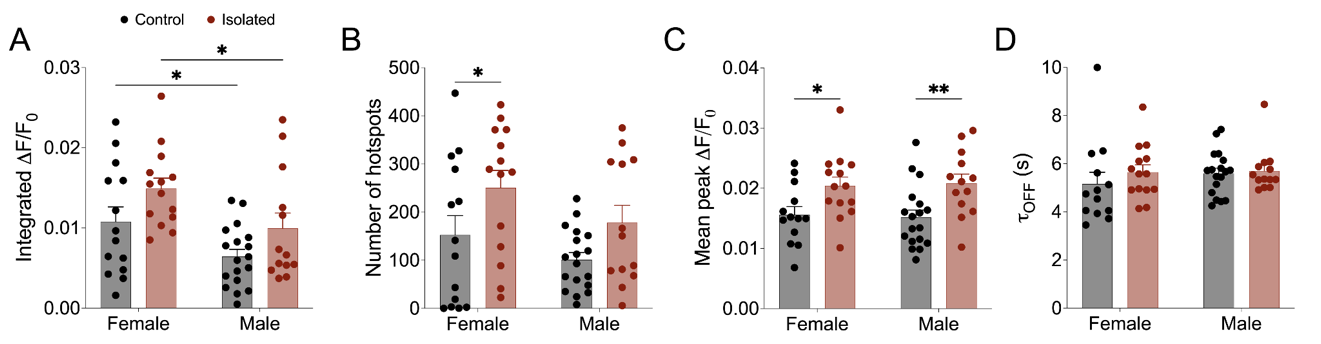

Fig. S4. Sex comparison in male and female voles for parameters from nIRCat imaging (mate relationship only). (*A*) Integrated ΔF/F_0_ in isolated (red) and control group (black) of female voles. (*B*) The number of hotspots identified in the field of view. (*C*) ΔF/F_0_ averaged over identified hotspots. (*D*) ΔF/F_0_ signal decay constant _OFF_ averaged over identified hotspots. n = 14 slices (6 animals) for control and n = 14 slices (6 animals) for female, and n = 18 slices (8 animals) for control and n = 13 slices (6 animals) for isolated groups for male. Non-significant comparisons are omitted for clarity. *  =  <.05, **  = <.01, *** <.001, **** < .0001 as determined by post-hoc Fisher’s LSD test. See Table S4 for details. Error bars denote the standard error of the mean.

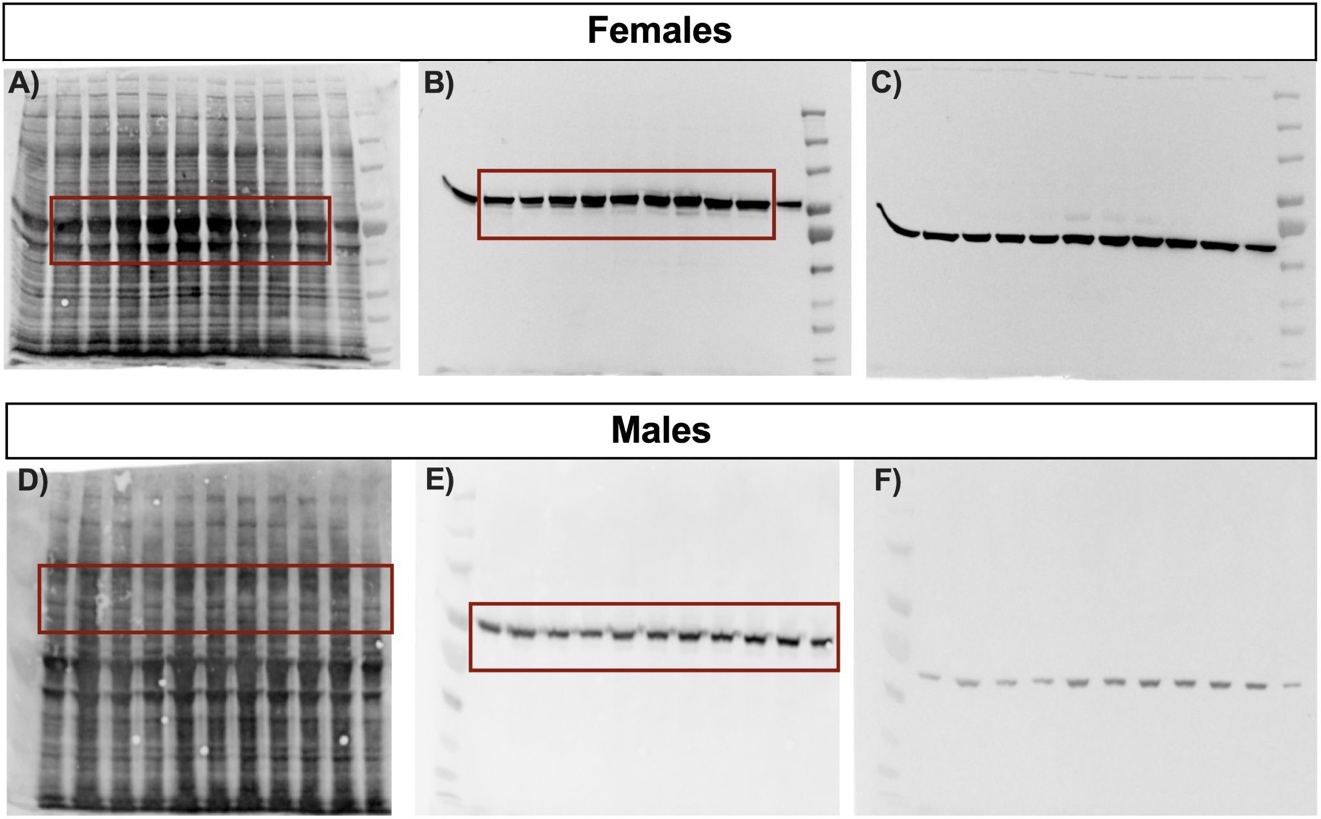

Fig. S5. Raw Western blot images with red box highlighting the bands shown in *Figure 2*. (*A-C*) Female Western blots showing (*A*) total protein, (*B*) tyrosine hydroxylase (TH), and (*C*) beta-actin concentration. Lanes 2-6 are control females, while lanes 1 and 7-10 are isolated animals. Lane 11 was not quantified in this experiment and served as an inter-assay control for a separate study. (*D-F*) Male Western blots showing (*D*) total protein, (*E*) TH, and (*F*) beta-actin concentration. Lanes 1-6 are control males, while lanes 7-11 are isolated males.
